# Standing genetic variation buffers field populations of *Zymoseptoria tritici* against seasonal and fungicide selection

**DOI:** 10.64898/2026.08.20.745885

**Authors:** Andrea Tobian Herreno, Pu Huang, Isabella Siepe, Remco Stam

## Abstract

- *Zymoseptoria tritici* is a fungal wheat pathogen whose exceptionally large effective population sizes and frequent sexual recombination enable rapid adaptation and the breakdown of disease control strategies, yet the relative contributions of demographic turnover and fungicide selection to within-season genomic change remain unresolved at the field scale.
- We analysed whole-genome sequences from five wheat fields sampled during epidemic progression, including paired untreated and fungicide-treated populations, to separate seasonal demographic change from fungicide effects on genome-wide diversity allowing us to compare minor allele frequency spectra and diversity statistics to disentangle these effectswithin individual fields.
- Field populations were locally differentiated yet nested within the broader European gene pool; within-season demographic turnover consistently shifted allele frequency spectra towards more shared, common alleles; whereas nucleotide diversity and adaptive potential remained largely unchanged.
- Seasonal demographic turnover accounted for most short-term genomic change, while fungicide effects were comparatively small, field-specific and acted primarily on pre-existing resistance alleles and standing genetic variation.
- Our results show that short-term adaptation is driven primarily by the redistribution rather than depletion of standing genetic variation, highlighting pathogen population biology as a key determinant of disease-control durability and emphasizing the value of population-informed genomic surveillance.

## Introduction

The durability of crop disease control is, in essence, an evolutionary question. Whether a fungicide or a resistance gene holds for a few seasons or many is determined less by the compound efficiency or resistance gene itself than by the evolutionary potential of the pathogen population it meets: its effective population size, its rate of recombination, and above all, the reservoir of heritable variation already segregating when selection begins (McDonald & Linde, 2002; Croll & McDonald, 2017; Stam & McDonald, 2018; Garnault *et al*., 2021). For fungal crop pathogens with large population sizes, adaptation typically proceeds from this standing genetic variation rather than from new mutation (Barrett & Schluter, 2008): the alleles that confer resistance are usually present at low frequency in advance, and selection acts by shifting their frequencies rather than by waiting for them to arise. A large effective population, recombining and fast-turning-over, is therefore both highly adaptable and difficult to exhaust because single-target control redistributes its reservoir without emptying it (McDonald *et al*., 2022). This reframes management as a population-genetics problem: to anticipate adaptation rather than record it once control has failed, we must characterise how much variation a field population carries, how it is structured, and how the selection we impose reshapes it. Crop diseases already threaten global food security, a burden expected to grow as climate change widens pathogen infection opportunities (Smith *et al*., 2020; Campbell *et al*., 2023; Singh *et al*., 2023), which makes this predictive knowledge more pressing.

Standing variation is informative as well as adaptive. In a large, recombining population that has not been recently bottlenecked, the site-frequency spectrum is dominated by rare, largely neutral alleles (Tajima, 1989), and summary statistics such as Tajima’s D and the minor-allele-frequency spectrum record this demographic history and flag departures consistent with selection (Croll & McDonald, 2017; Feurtey *et al*., 2023). The depth of this reservoir sets a ceiling on how fast a population can respond: the more independent genomes it maintains, the more low-frequency variants are available for selection to draw upon (Barrett & Schluter, 2008; McDonald *et al*., 2022). Reading these signals, and therefore judging durability, depends on sampling the underlying variation without bias and without over-representing clones, which places sampling design and depth at the base of any such assessment (McDonald & Mundt, 2016; Harrison *et al*., 2024).

*Zymoseptoria tritici* (Zt), an ascomycete hemibiotroph, causes Septoria tritici blotch (STB), a major constraint on wheat production. It reproduces asexually via pycnidia during the season and sexually via pseudothecia at season’s end; this mixed strategy, combined with a polycyclic infection cycle, long-distance ascospore dispersal and persistence in resistant structures such as chlamydospores, drives rapid adaptation and continual genotype turnover (Suffert *et al*., 2011; McDonald *et al*., 2015; Brown *et al*., 2015; Suffert & Thompson, 2018; Badet *et al*., 2020; Fagundes *et al*., 2020; Feurtey *et al*., 2023; Amezrou *et al*., 2023). Inoculum persisting in crop residues and soil further complicate disease management (Garnault *et al*., 2021; Hellin *et al*., 2021; Karisto *et al*., 2022; Klink *et al*., 2022).

Zt populations are gaining resistance to widely used fungicides while adapting to a changing climate, including rising temperature and radiation. European surveillance shows widespread resistance to demethylation inhibitors (DMIs) and succinate dehydrogenase inhibitors (SDHIs): resistance alleles in CYP51, the primary DMI target, are now common, with variants like I381V at high frequency across countries, and SDHI resistance is also rising (Hellin *et al*., 2021; Talas *et al*., 2026), indicating strong, ongoing selection with implications for the durability of chemical control (Hellin *et al*., 2021; Jørgensen *et al*., 2022; Kildea *et al*., 2025). Crucially, as mentioned above, these resistance alleles are recurrent and often pre-existing across countries rather than arising anew at each site, which is the field-scale expression of adaptation from standing variation (Garnault *et al*., 2021). Local environment modulates these dynamics (Suffert & Sache, 2011; Morais *et al*., 2015; Suffert *et al*., 2015) The number of infection windows and the latent period varies with temperature and moisture (mean approximately 20.35 days in northern Germany, shortening by approximately 0.95 days per 1 °C) (Henze *et al*., 2007), and decision-support systems exploiting such parameters can reduce spray frequency without losing control over disease severity, highlighting multiple opportunities for the fungus to infect tissues in varying stages of the wheat senescence and epidemic and the importance of functional fungicides (Burke & Dunne, 2008).

At the molecular level, virulence is shaped by effector diversity and its regulation. The IPO323 reference genome spans 39.73 Mb across 21 chromosomes (13 core, 8 accessory) (Goodwin *et al*., 2011; Lapalu *et al*., 2025), the accessory set showing frequent presence-absence variation and rapid loss under stress (Schotanus *et al*., 2015; Plissonneau *et al*., 2018; Badet & Croll, 2020). It encodes hundreds of candidate effectors, many highly variable through mutation, deletion and transposable-element activity (Amezrou *et al*., 2023). The well-characterised examples act in the biotrophic and asymptomatic phases with tightly timed, spatially restricted expression: *AvrStb6* is recognised at the stomata during penetration, while *Mg3LysM* and the quantitative *Avr3D1* and *AvrStb9* maintain the symptomless phase. Others peak at the biotrophic-to-necrotrophic transition that cell-death-inducing effectors and cell-wall-degrading enzymes then drive (Rudd *et al*., 2015; Yang *et al*., 2015; Meile *et al*., 2018, 2023, 2024; Thynne *et al*., 2024; Alassimone *et al*., 2024). The interplay between effector diversity and host recognition is crucial for resistance durability in wheat: 23 resistance (Stb) loci have been mapped but only three cloned (Brown *et al*., 2015; Saintenac *et al*., 2018; Battache *et al*., 2024; Hafeez *et al*., 2025), and effectors such as *Avr3D1* and *AvrStb9* already segregate as multiple recognition-evading alleles available to immediate selection (Kettles & Kanyuka, 2016; Saintenac *et al*., 2018, 2021; Battache *et al*., 2022, 2024; Suffert *et al*., 2024).

Zt populations have large effective population sizes and extensive variation, sustained by recombination, accessory chromosomes and transposable elements (Croll & McDonald, 2017; Dutta *et al*., 2021b; Singh *et al*., 2021; Hartmann *et al*., 2021). This is precisely what keeps the standing reservoir deep and continually replenished, so that most segregating variation is neutral and available rather than spent. Population-genomic analyses show clear global structure, with the highest diversity in the Middle East and North Africa near wheat’s centre of origin (Feurtey *et al*., 2023), and fine-scale studies reveal substantial local diversity, private alleles and recombination-driven structure (Siah *et al*., 2018; Chedli *et al*., 2022). Because genetically distinct isolates co-occur within single leaf layers, resolving within-field diversity without over-representing clones requires dense, hierarchical sampling (Hassine *et al*., 2019; Orellana-Torrejon *et al*., 2022a,b; Karisto *et al*., 2022; Tobian Herreno *et al*., 2025); such genotypic variation in turn influences virulence, dispersal and disease progression in host- and environment-dependent ways (Hartmann & Croll, 2017; Lorrain *et al*., 2024; Stapley *et al*., 2025; Talas *et al*., 2026).

Even so, field-scale population structure across European landscapes remains incompletely resolved, and the co-occurrence of clonal and sexually derived isolates raises questions about how dispersal, recombination and selection shape local epidemic dynamics (Hassine *et al*., 2019; Karisto *et al*., 2022; Bernasconi *et al*., 2022; McDonald *et al*., 2022; Tobian Herreno *et al*., 2025). We test whether a season of epidemic growth and a single azole application restructure field populations of *Z. tritici* by depleting standing genetic variation or by redistributing existing diversity. To do so, we characterised within-field population structure and genome-wide diversity across five fields in northern Germany over time, and used paired untreated and fungicide-treated populations to disentangle seasonal demographic change from fungicide effects. We further examined whether these shifts extend to effector and fungicide-resistance loci.

## Materials and methods

### Collection data and sampling

We surveyed five experimental winter wheat fields (cultivar Chevignon) across Schleswig-Holstein, Germany, in summer 2023: Dollerup and Futterkamp on the Baltic coast, Kating and Kollmar on th e North Sea coast, and Rade inland. Each field was sampled along two exponential transects in the outer lanes, at a centre point and 2, 8 and 32 m to either side (Supplementary Figure 2. A), following a BASF SE Group Research design. Two symptomatic leaves per plant were collected at growth stage (GS) 30 to 31 (stem elongation); a late-season collection at the same GPS points (GS 59 to 61, end of heading) sampled conventionally treated plants (Score; difenoconazole-) at all fields (approx. 1 week before sampling) and paired untreated plants from the same locations (protected through plastic cover at the days of treatment) at Dollerup and Kollmar. Single-pycnidium cirri were isolated on water agar, grown on malt-yeast agar with kanamycin (7 d, 18 to 20°C), and genomic DNA was extracted with the Mag-Bind Blood & Tissue HDQ 96 kit; Illumina short-read sequencing was performed in-house at BASF SE. Published UK (ENA PRJEB81422; (Tobian Herreno *et al*., 2025)) and Swiss and US (NCBI PRJNA596434; (Singh *et al*., 2021)) fields were reanalysed in parallel. German sequence data are deposited under ENA PRJEB[confirm]. All analyses used the IPO323 reference (Goodwin *et al*., 2011; Lapalu *et al*., 2025).

### Alignment and Variant Calling

Reads were aligned to IPO323 with BWA-MEM v0.7.17 (Li, 2013), sorted and indexed with SAMtools v1.17 (Li *et al*., 2009) and validated with Picard (GATK v4.0.5.1) (Poplin *et al*., 2017); isolates with core-chromosome depth below 10 were discarded. Following the standard variant calling pipeline for *Z. tritici* (Singh *et al*., 2021; Feurtey *et al*., 2023; Tobian Herreno *et al*., 2025), multi-sample genotypes were called per field with ploidy 1 with GATK HaplotypeCaller, CombineGVCFs and GenotypeGVCFs (maximum two alternate alleles, six genotypes per site) and hard-filtered with VariantFiltration (MQ < 20, QD < 2, QUAL < 30, SOR > 3, FS > 60, DP < 10, ReadPosRankSum and MQRankSum outside ±2). Biallelic SNPs were retained with VCFtools v0.1.16 (Danecek *et al*., 2011) at over 80% genotyping rate (--max-missing 0.8) and minor allele count of at least 1; MAF and per-site missingness were extracted with VCFtools and PLINK (Purce **l** *et al*., 2007). Accessory-chromosome presence or absence was called per isolate with CNVkit v0.9.10 (Talevich *et al*., 2016) in whole-genome mode against a flat pooled reference (haploid, --ploidy 1), scoring the length-weighted mean log2 ratio per chromosome as loss (< −0.5), partial loss (−0.5 to −0.2), neutral, partial gain (0.2 to 0.5) or gain (> 0.5). Copy number was compared between fields (Kruskal-Wallis) and between treatments within a field (two-sided Wilcoxon rank-sum), Benjamini-Hochberg corrected across chromosomes.

### PCA analysis and Admixture

Core-chromosome SNPs (chromosomes 1 to 13) from all eight fields were filtered to 5% MAF and thinned to one SNP per 10 kb. PCA was run on the GDS-converted VCF with SNPRelate::snpgdsPCA v1.38.1 (Zheng *et al*., 2012). Admixture was estimated with ADMIXTURE v1.3.0 (Alexander *et al*., 2009) on about 42 to 60 early-time point isolates per field, LD-pruned with PLINK --indep-pairwise (50-SNP window, 10-SNP step, VIF 0.2) and re-filtered to 5% MAF, at K = 1 to 15 with 10 replicates per K (Feurtey et al. 2023). Per-field PCA used the 15 isolates closest to the field centroid as representative.

### Diversity statistics

All statistics were haploid and restricted to the core chromosomes. Per field and tre atment we computed π, Watterson’s θ, Tajima’s D, Fu and Li’s D* and F*, and the singleton fraction with scikit-allel v1.3.7 (Miles *et al*., 2024), scored per whole chromosome and in 10 kb windows of at least 10 SNPs. D* and F* were computed in their folded, no-outgroup form (Fu & Li, 1993) from minor-allele singletons, an approximation appropriate to intraspecific data. Windows were treated as independent units on the basis of rapid LD decay. Because the singleton class is defined by a threshold of 1/n, the raw singleton fraction, D* and F* are not comparable between groups of unequal sample size, and our fungicide collections carry more isolates than their untreated controls in both paired fields (Dollerup 54 versus 40; Kollmar 60 versus 51). For all between-group comparisons the folded spectra were therefore down-projectedhypergeometrica **l**y to a common 34 isolates, the size of the smallest German collection, and the singleton fraction, D* and F* recomputed from the projected spectra (Supplementary Table 8b). Values at native n are reported alongside for comparison. Distributions were compared between groups with Kruskal-Wallis tests; where the same windows were available at both time points, the more conservative unmatched test is reported alongside a paired Wilcoxon signed-rank test on the per-window difference (Supplementary Tables 5c, 6, 8).

Synonymous and non-synonymous SNPs were annotated with SnpEff v5.2a (Cingolani *et al*., 2012) against the IPO323 gene models (Lapalu *et al*., 2025). Per field and treatment, a pN/pS ratio was calculated as the count of non-synonymous (missense, nonsense, frameshift and other protein-altering) over synonymous SNPs, each counted once per isolate carrying the alternative allele, scored per chromosome and in 10 kb windows, with windows of fewer than five coding SNPs excluded. For the two fields regular fungicide treated and additional untreated collections (Dollerup and Kollmar), per-window statistics were matched by window and the fungicide-minus-untreated difference tested against zero with Wilcoxon signed-rank tests, Benjamini-Hochberg corrected across statistics.

Within-field pairwise distances were the relative Hamming distance over jointly called sites; clonal groups were clusters below 0.01 relative distance (Singh *et al*., 2021). Isolation by distance was tested per group with a Mantel test (Spearman’s rank, 9,999 permutations, Benjamini-Hochberg) between the relative-Hamming and a haversine geographic matrix, run on the full isolate set and again after collapsing each clonal cluster to one representative.

Distribution of haplotypes of *AvrStb6*, *AvrStb9*, *Avr3D1, ZtIPO323_017790, ZtIPO323_019470, ZtIPO323_106990, ZtIPO323_117500, ZtIPO323_119610, and ZtIPO323_122170* (Rudd *et al*., 2015; Amezrou *et al*., 2023; Lapalu *et al*., 2025) were drawn as minimum spanning networks with poppr (Kamvar *et al*., 2014), and multilocus-genotype distributions compared between fields and between treatments with Fisher’s exact tests and Benjamini-Hochberg correction.

### Allele-frequency change quantification

Folded MAF spectra of the early samples were compared between fields as a KS distance, calibrated against a noise floor of KS distances between UK subsamples and by pooled-label permutation (500 permutations, add-one p, Benjamini-Hochberg within the German-German and German-UK families); a Kruskal-Wallis omnibus is reported. For the paired fields, per-site ΔAF (treated minus untreated) was computed with scikit-allel, with an empirical p from 200 same-timepoint splits of the untreated population; loci in the top 5% of absolute change (95th percentile) were called significant, and per-chromosome ΔAF distributions compared with KS tests (Benjamini-Hochberg across the 13 chromosomes). Each field’s observed 95th-percentile ΔAF in 10 kb windows was compared with a split-half null from randomly halving the early population, and with split-half nulls from subsampled Swiss and UK isolates. Folded site-frequency spectra were projected to a common n = 33 haploid genomes with the hypergeometric estimator and normalised per frequency class. Per-isolate CYP51 coding sequenceswere reconstructedas a haploid consensus against IPO323 with low-coverage positions masked, translated, and compared with the reference at known azole-resistance positions (Garnault *et al*., 2021; Hellin *et al*., 2021; Kildea *et al*., 2025); amino-acid frequencies were compared between paired untreated and fungicide samples by Fisher’s exact test with Benjamini-Hochberg correction. Deletions at Y459/G460 were not assessed, as indels were excluded upstream.

### Identifying selective sweeps between treatments and climate differences

Selective sweeps were detected with SweepFinder2 v1.0 (DeGiorgio *et al*., 2015) per field and treatment on a 10 kb grid across the core chromosomes; treatments were compared pairwise per chromosome with two-sample KS tests (Benjamini-Hochberg). Monthly bioclimatic variables (WorldClim v2.1; (Fick & Hijmans, 2017)) were sampled at field GPS points with the terra package and compared among fields with Kruskal-Wallis and paired Wilcoxon tests (Benjamini-Hochberg). Figures were produced with ggplot2 v3.5.0 in R v4.2.1, and matplotlib and seaborn in Python.

### Dataset resources

The UK field sequence data is available at Bioproject PRJEB81422. The sequencing data for the Swiss and American fields are available in NCBI in BioProject PRJNA596434 (Singh *et al*., 2021). We used the reference genome of IPO323 for variant calling (Goodwin *et al*., 2011; Lapalu *et al*., 2025), which is available at https://mycocosm.jgi.doe.gov/Zymtr1/Zymtr1.info.html

All the scripts present in this study can be found at https://github.com/PHYTOPatCAU/ZymoFieldDiv_Germany.

## Results

### The German fields are densely sampled and fall within the European population

To assess within field dynamics of *Zt,* we surveyed five experimental wheat fields across Schleswig-Holstein: Dollerup and Futterkamp on the Baltic coast, Kating and Kollmar on the North Sea coast, and Rade inland. Temperature ranges were similar across fields, but precipitation differed (Kruskal-Wallis p = 0.002; Supplementary Table 1). In total, we genotyped 631 isolates, a mean of 126 per field, at two time points: an early, pre-treatment collection (GS30 to 31; 247 isolates, mean 49 per field) and a late collection (GS59 to 61; 384 isolates) of difenoconazole-treated plants at all fields and paired untreated plants at Dollerup and Kollmar (Supplementary Table 2). Sequencing depth averaged 41x, comparable to the published UK, Swiss, and US datasets (47x, 69x, and 32x), and supported high quality genotyping: mean call rate above 90% per field and, after stringent filtering (per-site missingness below 20%), high-confidence genome-wide SNPs (Supplementary Figure S1A, B) (Hartmann *et al*., 2018; Singh *et al*., 2021; Feurtey *et al*., 2023).

To assess the genetic background of the isolates we performed an admixture analysis for all early-time point isolates and subsampled international fields. At optimal K, all newly sampled isolates show similar ancestry as the previously sampled UK and CH fields. The PCA, resolvedtwo genetic clusters, one European and one from the US, with the German samples clearly within the European cluster (Figure 1A, B; Supplementary Figure S2B; Supplementary Table 3). This is consistent with previous work (Hartmann *et al*., 2018; Feurtey *et al*., 2023) and confirms that our German populations are included in the main European *Z. tritici* population. For the following analyses we therefore kept the UK and German fields, all sequenced between 2023 and 2025 with similar depth and technology.

**Figure 1.**
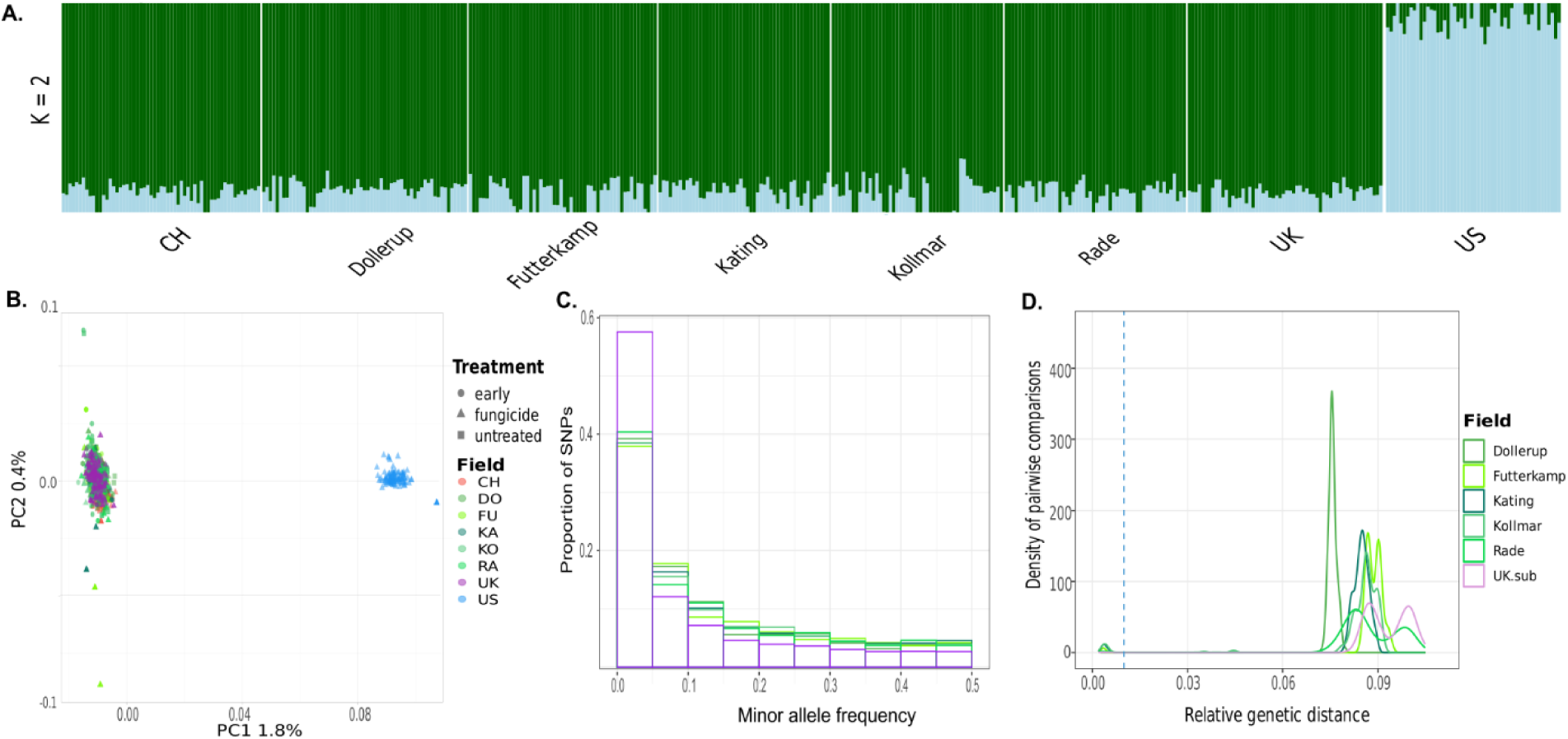
Baseline population structure and sampling locations show the European population with differences on their site frequency spectrum and genetic distance distribution. **A.** Admixture of early German isolates and subsampled external isolates from Switzerland (CH), the United Kingdom (UK) and the United States (US) (n approximately 60 per field) using core-chromosome SNPs (MAF 5%, thinned to one SNP per 10 kb) at K = 2; each bar is one isolate, coloured by ancestry proportion. **B.** Whole-dataset PCA on core-chromosome SNPs (MAF 5%). The first component separates the US isolates from the European cluster. **C.** Whole-genome folded MAF spectra for the selected European fields as the proportion of SNPs per MAF bin, each normalised to its own SNP total. Early samples (GS 30 to 31, elongation), n approximately 42 to 60 per field; the UK field is a late (GS 83), fungicide-treated subsample (n approximately 60) retained for comparison of the starting variation present at each field, where each distribution differs between most fields except Kollmar-Rade (Only comparison with Permutation p value with Benjamini-Hochberg correction p > 0.05). GS, growth stage; MAF, minor allele frequency; SNP, single -nucleotide polymorphism. **D.** Whole-genome pairwise genetic-distance distributions per field (relative Hamming distance); the dashed light-blue line marks the 1% clonality threshold.

### The Minor allele frequency spectra and genetic distance distribution of the fields are different at the early time point

We next asked whether the five field populations shared common genomic characteristics or instead represented distinct local populations. Before comparing fields, we confirmed that sampling 60 isolates per field provided sufficient power for population genomic inference. Independent random subsamples of 60 isolates from the UK population produced highly consistent minor allele frequency (MAF) spectra and population genetic summary statistics (π, θ and Tajima’s D; Supplementary Table. 4; Supplementary Table 5b), demonstrating that the observed differences among German fields reflect biological variation rather than sampling effects.

The minor allele frequency spectra (MAF) showed a typical the left-skewed shape similar as seen in previous studies (Singh *et al*., 2021; Tobian Herreno *et al*., 2025), with most variants present at low frequency. The German fields were, however, less enriched in singletons than the UK (Supplementary Table 5c). The UK sample is late and fungicide-treated, so this difference is not a seasonal effect: between-population differences in the depth of the rare tail exceed the within -season shift we measure below.

Each field showed a unique variant distribution except for the Kollmar-Rade comparison (Permutation p < 0.05; Figure 1C; Supplementary Table 4). Genetic distances broadly reproduced the earlier study (Tobian Herreno *et al*., 2025): most pairwise comparisons were near-equidistant, with a few clonal clusters below the 1% core-genome dissimilarity threshold (Figure 1D). Kollmar, Kating, Rade and Futterkamp showed additional peaks beyond that threshold, hinting at weak within-field structure, though this was not significant in a field-level PCA (Supplementary Figure S3A). Overall, this confirms that the five sampled fields lie firmly within the European population and even with the maximum distance between any two fields being approximately 150 km, each field has their own MAF and genetic distance distributions.

### Epidemic progression diversity the same way in every field

Looking at their nucleotide diversity statistics, at the early time point the German fields did not differ in nucleotide diversity (π): every pairwise comparison between German fields was non-significant (median π ≈ 0.0102 in each), and π separated the German fields only from the UK (pairwise Wilcoxon rank-sum with Benjamini-Hochberg; Supplementary Table 5; Figure 2A). The fields differed instead in the shape of the rare tail. Watterson’s theta also changes between time points, decreasing from the early to the late time point (Figure 2B; Supplementary Table 6). Tajima’s D also differed (p < 0.05; Figure 2C; Supplementary Table 5; Table 5c), with Futterkamp and Kollmar the least negative and Kating and Rade the most, a grouping reproduced by Fu and Li’s D* and the projected singleton fraction. Medians stayed negative, consistent with European populations (Singh *et al*., 2021; Tobian Herreno *et al*., 2025), but the German fields sat more positive than the UK, reflecting their lower singleton richness. The mean MAF also shifted toward the more positive values, displaying the depletion of lower frequency alle les (Figure 2 D), consistent with the depletion observed in the normalised n per field singleton fraction between time points (Figure 2 E; Supplementary Table 6). Accessory-chromosome copy number differed on chromosomes 18 and 21 between fields (Kruskal-Wallis on the per-chromosome mean log2 ratio with Benjamini-Hochberg, p < 0.05; Supplementary Figure S3B).

**Figure 2.**
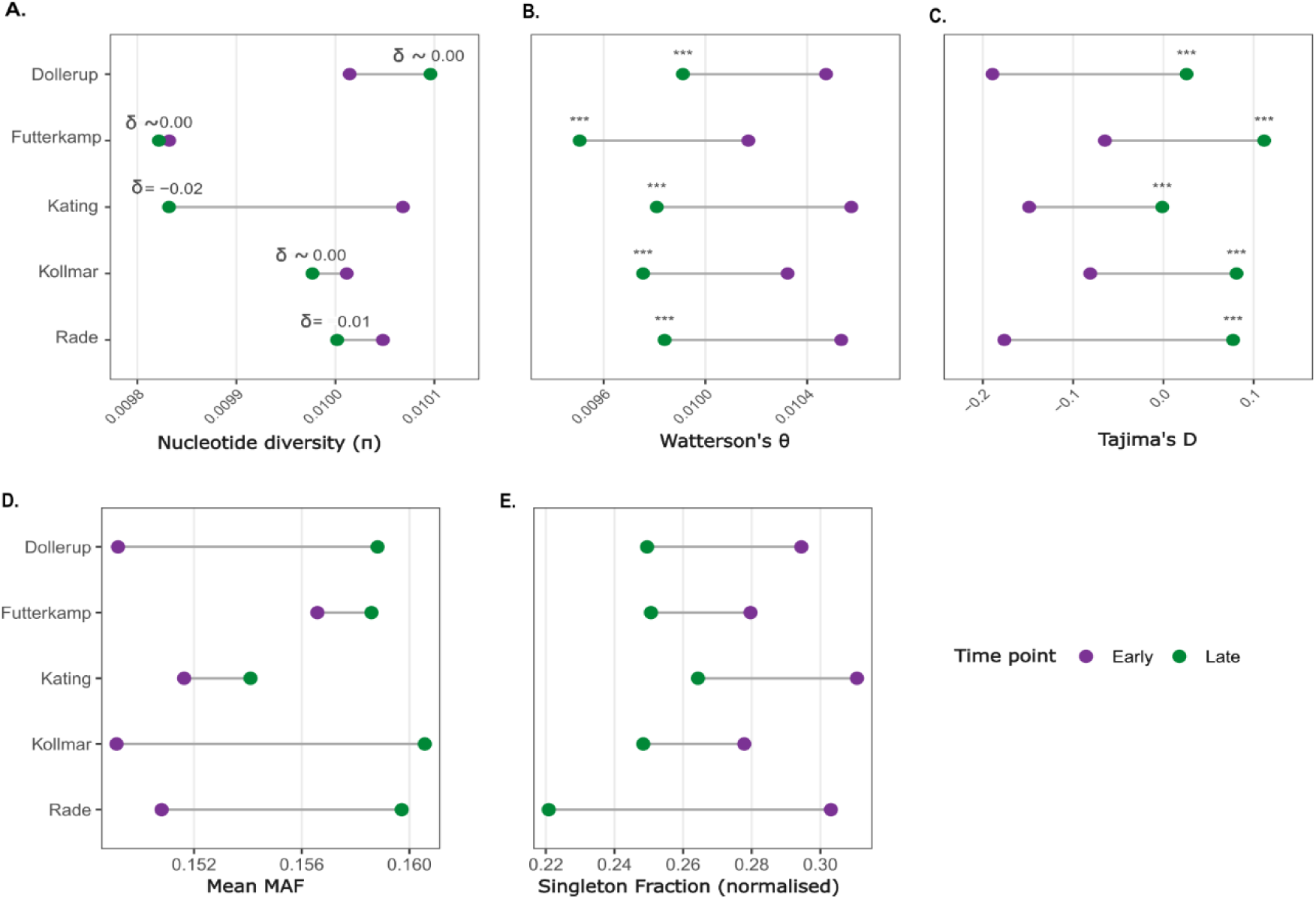
Epidemic progression shifts the field populations towards a larger common-variant fraction and a thinned rare tail. **A.** Nucleotide diversity (π) per field. π does not shift significantly across timepoints in any field; the common-variant pool is preserved through the epidemic season. **B.** Watterson’s θ per field. θ falls from early to late in every field, the counterpart to the Tajima’s D rise in (C): together they mark a loss of low-frequency variants without a change in overall diversity. **C.** Tajima’s D per field. D rises from early to late in every field as rare alleles are trimmed. **D.** Mean minor allele frequency per field. Mean MAF rises from early to late in Dollerup, Futterkamp, Kating, Kollmar and Rade as the spectrum shifts away from its rare tail. **E.** Singleton fraction, computed from folded MAF spectra projected to a common isolate count (n = 34) to remove sample-size confounding. Singletons decline from early to late in every field, quantifying the rare-variant depletion inferred from θ and Tajima’s D. Statistics computed in 10 kb windows genome-wide on the core chromosomes. Panels A to E are dumbbell plots showing the median per field at each timepoint, joined by a rule spanning the within-field range. Between-timepoint tests are Wilcoxon signed-rank tests on the per-window difference, with Benjamini-Hochberg correction across the five fields per statistic (Supplementary Table 6); omnibus Kruskal-Wallis tests on the window distributions are reported in Supplementary Table 5c. Panel A reports Cliff’s δ per field in place of a star because π does not differ between time points in any field (Kruskal-Wallis p > 0.05; Supplementary Table 5c). Symbols: dot colour indicates timepoint (Early, Late). GS, growth stage; MAF, minor allele frequency; SNP, single-nucleotide polymorphism.

From this early baseline, every German field shifted the same way by the later time point with a change larger than the sampling variation of the UK and CH reference metapopulations (bootstrap 95% intervals; Supplementary Table 5b). Watterson’s θ fell in all five fields (paired per-window Wilcoxon signed-rank, matched by window; Benjamini-Hochberg across fields within each statistic, all p adjusted < 0.001; Supplementary Table 6), while Tajima’s D rose and moved from strongly negative towards zero (median shift +0.16 to +0.24). π stayed essentially flat: median shifts were at most 2 × 10⁻⁴ in every field, two orders of magnitude smaller than the shift in θ and inconsistent in direction, so the common -variant pool is unmoved and the change is not significant (Kruskal-Wallis p > 0.05; Supplementary Table 5c). This concordant pattern, θ down and Tajima’s D up with π held, is the signature of a seasonal purge of low - frequency variants, and its direction was uniform across all five fields.

### Timing of sampling drives changes in MAF and Tajima’s D and the observed rare-variant depletion

The analyses above describe the genomic changes occurring over the course of a growing season under conventional disease management. However, the comparison between early and late fungicide -treated populations combines two processes: seasonal demographic change during epidemic progression and the effect of fungicide application itself. To disentangle these components, untreated control plots were additionally sampled at the late time point in two fields (Dollerup and Kollmar). This allowed the overall seasonal shift to be partitioned into a demographic component (early to late untreated) and a fungicide component (late untreated to late treated), where the latter isolates the effect of fungicide because the two late populations differ only in treatment and were collected on the same day. Across the conventionally managed fields (early to late treated), lowest-frequency SNPs were depleted in both fields (Kolmogorov-Smirnov p < 0.05; Figure 3 C, F; Supplementary Table 7). Nucleotide diversity (π) did not change across the three treatments; early, late untreated and late, in either field (Kruskal-Wallis p > 0.05; Figure 3A; Supplementary Table 5c), whereas Watterson’s θ and Tajima’s D, did differ across samples (Kruskal-Wallis p < 0.05; Figure 3B, C, Supplementary Table 5c). Mean MAF shifted accordingly (Figure 3 E) toward a bigger value. - This is consistent with a redistribution of rare variants rather than a loss of overall diversity. From the early to the late untreated populations, Tajima’s D increased in both fields, moving from strongly negative values towards zero (Dollerup −0.317 to −0.016; Kollmar −0.242 to −0.159). While Watterson’s θ declined. The ratio of non-synonymous to synonymous mutations showed no changes throughout the season (Kruskal-Wallis p > 0.05 between treatments within fields; Figure 3 D, Supplementary Table 8).

**Figure 3.**
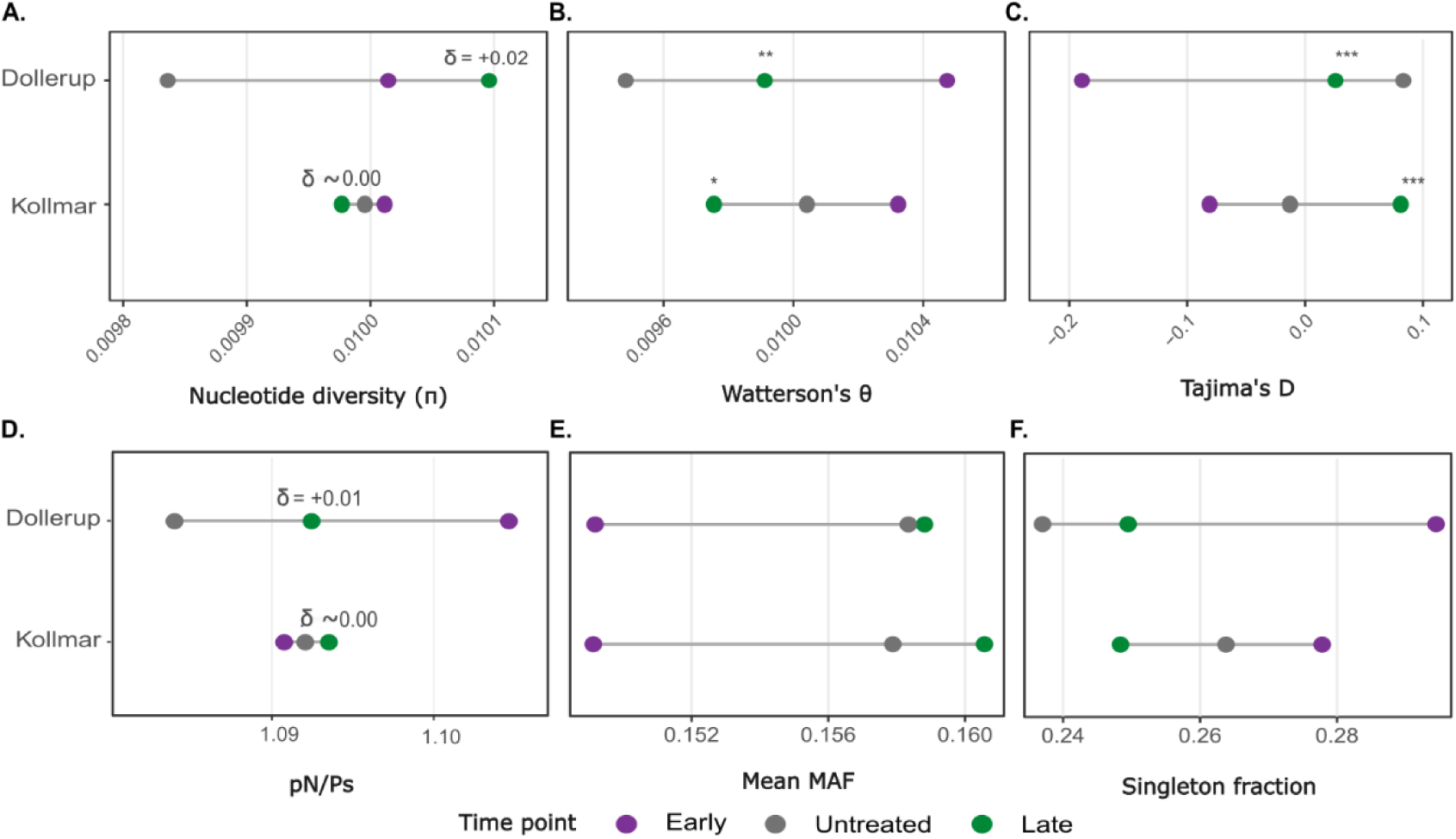
Season reshapes the rare tail while the common-variant pool holds, and only the fungicide step diverges between fields. The paired fields Dollerup and Kollmar, each sampled early (GS 30 to 31), late-untreated and late-fungicide (both GS 59 to 61, end of heading); the two late arms were collected on the same day and differ only in treatment. **A.** Nucleotide diversity (π) across time points; the π distributions do not differ within either field (Kruskal - Wallis p > 0.05; Supplementary Table 5c), so the common-variant pool is unmoved by season or spray. Cliff’s δ is reported in place of a significance star. **B.** Watterson’s θ across time points; θ differs between treatments in both fields (Kruskal-Wallis p < 0.05) and falls as the rare tail thins, the mirror of the Tajima’s D rise in (C): together they mark a loss of low-frequency variants rather than a change in overall diversity. **C.** Tajima’s D rises from early to late in both fields as rare alleles are trimmed. The two fields then differ at the spray step: Kollmar’s fungicide arm sits above its untreated control, Dollerup’s below it. **D.** pN/pS ratio per treatment; no difference between treatments (Kruskal-Wallis p > 0.05), placing the shift outside protein-altering variation and pointing to a demographic rather than a selective response. **E.** Mean minor allele frequency (MAF) per treatment. Mean MAF rises from early to late in both fields as the spectrum shifts away from rare variants, tracking the θ decline and Tajima’s D rise in (B, C). **F.** Singleton fraction, computed from folded MAF spectra projected to a common isolate count to remove sample -size confounding. Both fields carry fewer singletons late than early. Panels A to D are computed in 10 kb windows on the core chromosomes (chr 1 to 13); panels E and F are genome-wide. Between-treatment tests are Wilcoxon signed-rank on the per-window difference with Benjamini-Hochberg correction (*p < 0.05, **p < 0.01, ***p < 0.001; Supplementary Table 8); omnibus Kruskal-Wallis tests on the window distributions are in Supplementary Table 5c, and genome-wide spectrum statistics in Supplementary Table 8b. Cliff’s δ is reported where the omnibus is not significant. GS, growth stage; MAF, minor allele frequency; SNP, single -nucleotide polymorphism.

Comparing early with late-untreated first, both fields moved the same way; only the magnitude differed. Tajima’s D rose in each (Dollerup −0.317 to −0.016; Kollmar −0.242 to −0.159), Watterson’s θ fell in each, and the projected singleton fraction fell in each (Dollerup 0.294 to 0.237; Kollmar 0.278 to 0.264). Dollerup’s seasonal step was three to six times the size of Kollmar’s on every statistic (Figure 3B, C, F; Supplementary Table 8). The seasonal reshaping is therefore shared, and the fields have not yet diverged.

The untreated and fungicide-treated plants were collected at the same time, so this contrast compares two meta-populations that differ only in treatment. The two fields differ in the direction of that difference. In Kollmar the fungicide-treated population carries fewer low-frequency variants than its untreated control: the projected singleton fraction is lower (0.264 to 0.248) and Fu and Li’s D* higher (−0. 097 to +0.030). In Dollerup the difference runs the other way: the singleton fraction is higher (0.237 to 0.250) and D* lower (+0.123 to +0.020), with the excess confined to the singleton class, as the doubleton fraction fa **l**s (0.181 to 0.152; Supplementary Table 8b). Because singletons and doubletons move in opposite directions in Dollerup, Tajima’s D nets out to near zero for this contrast and the singleton -specific statistics resolve it more clearly.

The fungicide step is both smaller than the season and opposite in sign between the two fields: the shift in Fu and Li’s D* between the late arms is −0.10 in Dollerup and +0.13 in Kollmar, against shared early-to-late shifts of +0.37 and +0.24 (Supplementary Table 8b). Fungicide therefore modulates the trajectory the epidemic was already on, in a field-specific direction, rather than driving a change of its own scale.

If the seasonal shift reflects local expansion of a limited number of successful genotypes plus their local propagation, then nearby-sampled genotypes should be more similar late in the epidemic than distant ones. Clone-corrected isolation by distance (IBD) was absent early in every field (the only early signal, in Kollmar, vanished after clone collapse) but retained by several late samples (Dollerup untreated Mantel r = 0.56; Futterkamp late-treated r = 0.29; Kating late-treated r = 0.21; Kollmar marginal after correction, r = 0.222, BH p = 0.0585; Rade none; Supplementary Table 9). No consistent difference emerged between untreated and treated late populations, so fine-scale spatial structure develops during epidemic progression, not as a consequence of a fungicide application.

The same demographic process predicts a second signature: if late-season populations are dominated by descendants of a smaller pool of successful genotypes, accessory chromosomes, which are dispensable and stochastically lost, should show elevated loss under fungicide where the effective bottleneck deepens most (Habig et al., 2017; Möller et al., 2018). A directed early-to-fungicide Wilcoxon rank-sum flagged chromosomes 16 and 21 in Kollmar (BH p = 0.040 for both), and chromosome 16 across the pooled German fields (BH p = 0.020), with the same direction but no significance in Dollerup (BH p = 0.20; Supplementary Table 11a, b). Selective sweeps showed a related partial-overlap pattern: those detected early and between late treatments largely did not coincide, their distributions differed (per-chromosome Kolmogorov-Smirnov, BH p < 0.05; Supplementary Figure S6A, B; Supplementary Table 10), and in Kollmar post-treatment sweeps carried higher composite likelihood ratios in SweepFinder2. Together, IBD, accessory chromosome loss and sweep intensity point to localised rather than genome -wide responses, concentrated in the field where the seasonal purge deepened most under treatment.

### Allele-frequency composition shifts genome-wide across time points, while target-site variation is largely pre-existing

The observed changes in population genetic summary statistics imply widespread shifts in allele -frequency composition. We therefore asked whether these changes were distributed across the genome or concentrated at the CYP51 fungicide target. Across all three contrasts, loci in the top 5% of absolute allele-frequency change were distributedacross everycore chromosome and across the accessory genome, with no excess in the CYP51 window on chromosome 7 (Figure 4A, B; Supplementary Figure S5A), and the observed ΔAF distributions exceeded within-field sampling noise in all five fields (Supplementary Figure S2E, F). Per chromosome, the ΔAF distributions differed between treatments (Kolmogorov-Smirnov with Benjamini-Hochberg, P < 0.05; Supplementary Table 7), consistent with the loss of low-frequency variants captured by the diversity statistics rather than a single localised sweep. The change in allele-frequency composition through the season and under treatment was therefore population-wide and distributed across core and accessory chromosomes, rather than concentrated around the CYP51 window on chromosome 7. CYP51 showed extensive standing variation and carried resistance-associated substitutions at high frequency before treatment: I381V was fixed (proportion = 1) in every field and time point, and A379G, S524T and the multiallelic V136A/C already segregated at intermediate to high frequency at the early time point, in line with these alleles being long-established in the European CYP51 landscape (Huf *et al*., 2018; Hellin *et al*., 2021; Kildea *et al*., 2025) (Figure 4C; Supplementary Figure S5B; Supplementary Table 12). Only Kollmar showed a significant fungicide-associated shift in CYP51, an increase in S524T (Benjamini-Hochberg p = 0.019) and V136A and A379G marginally non-significant (p = 0.06); Dollerup showed no fungicide-specific change. Apparent fungicide-associated shifts in Futterkamp (V136A, S524T) and Kating (S524T) could not be separated from seasonal change, as those fields lacked an untreated control, and Y461H/S varied between time points but not between untreated and fungicide. Thus, azole application produced only local, field-specific changes in CYP51 against a background of extensive standing target-site variation.

**Figure 4.**
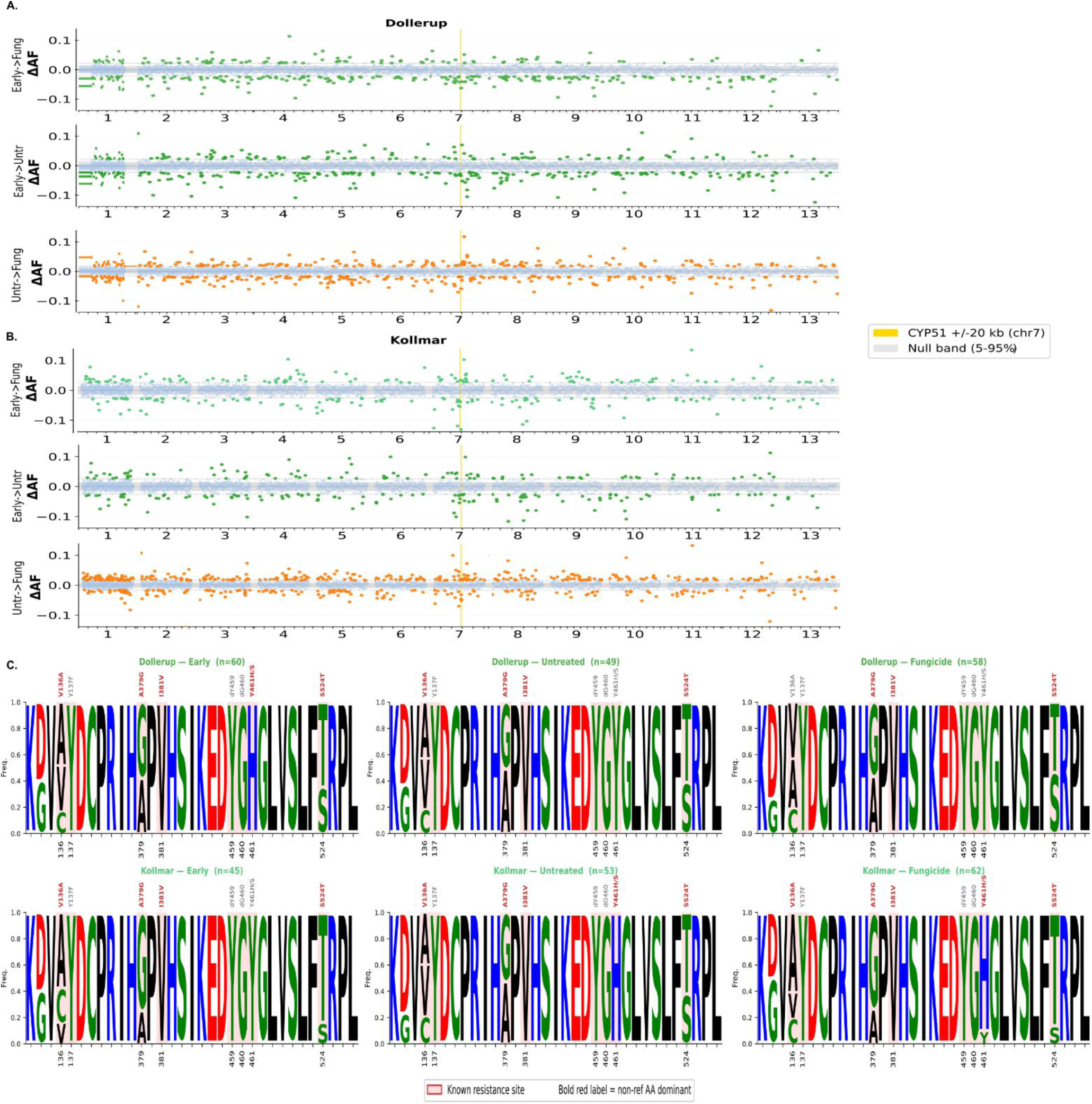
Allele-frequency change is genome-wide and not confined to the fungicide target, and CYP51 resistance is largely pre-existing. **A.** Per-site allele-frequency change in Dollerup for three contrasts: early to fungicide, early to untreated (seasonal), and untreated to fungicide. Loci in the top 5% of absolute allele-frequency change are coloured by contrast; the grey band is the 5th to 95th percentile of a permutation null built from same-timepoint splits of the untreated population. The CYP51 window is highlighted in yellow. Changed loci sit on every core chromosome rather than clustering at CYP51, marking the shift as a population-wide demographic reshuffling rather than a single localised sweep. **B.** Per-site allele-frequency change in Kollmar for the same three contrasts, analysed identically; the same dispersed, genome-wide pattern holds, confirming that neither the seasonal nor the spray step is driven by the fungicide target alone. **C.** CYP51 amino-acid motif logos for Dollerup and Kollmar at the early and late time point, untreated and treated. Literature-evidenced resistance sites are boxed in red; a bold red label marks positions where the non-reference allele dominates. Resistance substitutions are already dominant before treatment, so azole application perturbs CYP51 composition only locally against a background of high standing target-site variation.

### Demographic reshuffling, not loss, reshapes effector haplotype abundance within and between fields

We next asked whether the genome-wide demographic restructuring also extended to functionally important effector genes. We scored multilocus haplotypes (MLGs) for nine functionally characterised effectors, of which three (Avr3D1, AvrStb6 and AvrStb9) are shown in Figure 5A; the remaining six were tested but are not plotted (Supplementary Tables 13 and 14). Minimum spanning networks for the three shown are in Supplementary Figure S4. They show same demographic signal, a reshuffling of frequencies without loss of the underlying variants. Across all nine, no major private haplotypes separated the fields and a single haplotype dominated per field, though the identity of the dominant haplotype differed (Supplementary Table 13; Figure 5 A). At the early time point, haplotype composition differed significantly between fields for six of the nine effectors (Fisher’s exact with Benjamini-Hochberg, p ≤ 0.001; Supplementary Table 13); the remaining three did not differ between fields.

**Figure 5.**
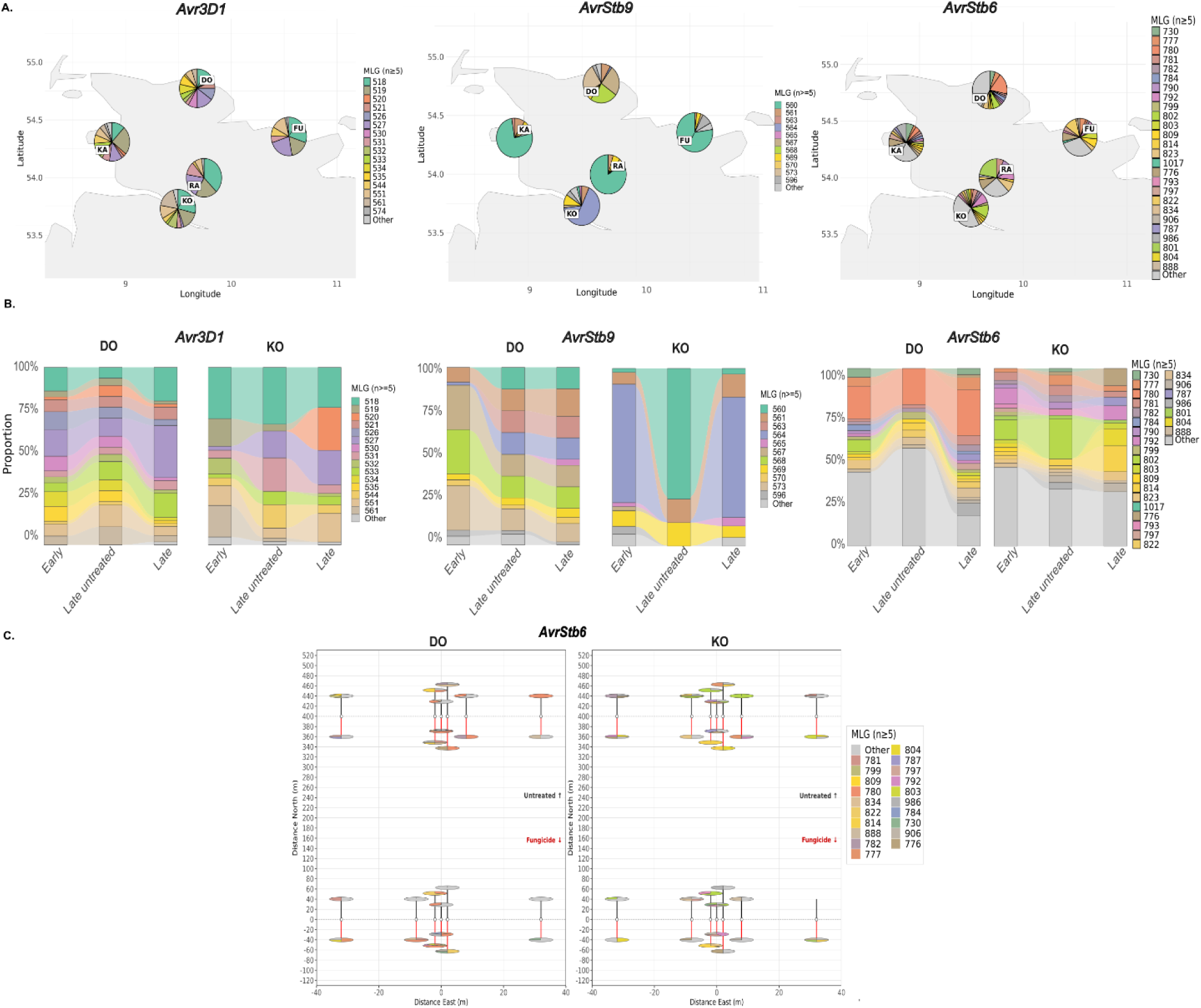
Effector haplotype dominance differs between fields at the outset and switches under treatment, without losing haplotypes. **A.** Distribution of the major multilocus genotypes (MLGs) of *AvrStb6, AvrStb9* and *Avr3D1* across the German fields, one pie chart per field at its sampling location, showing the frequency of each MLG (n ≥ 5) at the early time point. Each effector has a different dominant MLG in each field already at the early time point, and this between-field difference in composition is significant for all three genes (Fisher’s exact with Benjamini-Hochberg, p ≤ 0.001; Supplementary Table 13), so fields start from distinct haplotype balances rather than a shared one. **B.** Alluvial plot of MLG frequency shifts between early, late-untreated and late-fungicide in Dollerup and Kollmar. The dominant haplotype itself switches under treatment in Kollmar for all three effectors (Fisher’s exact with BH, p < 0.05), whereas in Dollerup only *AvrStb6* moves and only marginally (adjusted p = 0.051), with *AvrStb9* and *Avr3D1* unchanged (Supplementary Table 14). Ribbon widths reflect within-field proportions; no major haplotypes are gained or lost between treatments. **C.** Per-plant haplotype (MLG) composition of *AvrStb6* for Dollerup and Kollmar across the sampling points, one pie chart per plant, for both late -season treatments showing that the same composition change plays out across the field and that multiple haplotypes coexist within a single plant. MLG, multilocus genotype.

Between early and late sampling, haplotype frequencies shifted without gain or loss of major haplotypes, indicating a redistribution of existing variation (Figure 5B). These shifts were strongest in Kollmar, where six of the nine effectors changed significantly, including all three shown (Fisher’s exact with BH, p < 0.05). In Dollerup two of the nine changed (*ZtIPO323_017790 and ZtIPO323_117500*), while of the three shown only *AvrStb6* moved, and only marginally (adjusted p = 0.051; Supplementary Table 14). Fine-scale maps for Dollerup and Kollmar show the same composition change across sampled points and reveal multiple haplotypes within single plants (Figure 5C; Supplementary Figure S6C).

## Discussion

### Short-term evolution redistributes standing variation rather than depleting it

Our results show that short-term evolution in field populations is dominated by redistribution of standing genetic variation rather than its depletion. Across every level, from genome-wide allele frequencies to characterised effector and target-site loci, management and seasonality reshuffled pre-existing variation without depleting it. Standing diversity, not new mutation, is therefore the currency of short-term adaptation in this pathogen, and the selection episode we imposed shifted allele frequencies within a reservoir it did not exhaust. This is not peculiar to Zt: adaptation from standing variation is the rule across crop pathogens, seen in convergent azole resistance arising repeatedly from segregating CYP51 variants (McDonald *et al*., 2019), in effectors such as *Avr3D1* and *AvrStb9* that already segregate as multiple recognition-evading alleles (Meile *et al*., 2018, 2023; Amezrou *et al*., 2023), and in the standing-variation-fuelled host shifts of the pathogen *Cercospora beticola* (Chen *et al*., 2024; Taliadoros *et al*., 2025). What our paired design adds is the demonstration that a full season of epidemic growth and an azole application, the two forces expected to erode this reservoir, instead redistribute it.

The German populations formed part of the broader European *Z. tritici* population while remaining locally differentiated, consistent with previous observations (Hartmann *et al*., 2018; Feurtey *et al*., 2023; Tobian Herreno *et al*., 2025) This local differentiation likely reflects differences in the timing and composition of inoculum arriving in individual fields (Suffert *et al*., 2011; Suffert & Sache, 2011). Despite this, clonal clusters occur in every field but generated only weak intra-field genetic structure. This fits the pathogen’s epidemiology. Short-range splash dispersal seeds most local spread, while frequent sexual recombination and long-distance dispersal of ascospores reshuffles and redistributes genotypes, so even neighbouring isolates need not be related. Frequent recombination is expected to limit long-term clonal expansion by continually reshuffling genetic backgrounds, helping maintain the high standing diversity characteristic of this species. These observations further emphasise that local evolutionary dynamics are best captured by dense within-field sampling rather than sparse regional surveys (McDonald *et al*., 2022; Suffert *et al*., 2024).

### Standing genetic variation persists despite seasonal selection

Despite clear differences among fields, all European populations shared the same overarching genomic signature. Core-genome diversity (θ, Tajima’s D, Fu and Li’s D*) is significantly heterogeneous between fields at the early collection point, as expected for a mostly constraint free population (Tajima, 1989) (Figure 2B, C; Supplementary Table 5, 5c), while π is not, so the fields share a common-variant pool and differ only in the shape of the rare tail. Yet, all European fields share a negative Tajima’s D, an excess of low-frequency alleles (Tobian Herreno *et al*., 2025), without a collapse in π. This indicates no recent bottleneck and a large, recombining effective population size sustaining standing variation (Croll & McDonald, 2017; Hartmann & Croll, 2017; Hartmann *et al*., 2018; Feurtey *et al*., 2023), providing the material for rapid adaptation to a novel fungicide or resistance gene (Barrett & Schluter, 2008; Dutta *et al*., 2021a; McDonald *et al*., 2022). Importantly, neither seasonal epidemic progression nor fungicide application measurably depleted this. Nuclotide diversity (π) and the pN/pS ratio remained unchanged in the paired fields, whereas the shifts in θ and Tajima’s D were confined to the rare end of the allele frequency spectrum, again indicating redistribution of standing variation rather than widespread loss of functional diversity. (Henze *et al*., 2007; Burke & Dunne, 2008; Suffert *et al*., 2011; Morais *et al*., 2015), The absence of private effector haplotypes between fields and observations that haplotypes only shift in dominance suggests that local selection acts on a regionally shared reservoir maintained by recombination(Barrett & Schluter, 2008; Orellana-Torrejon *et al*., 2022a,b; Lorrain *et al*., 2024).

### Timing in the season, not spray, dominates short-term genomic change

An important implication is that field populations cannot be interpreted independently of their sampling context. Geography, epidemic stage and management are inevitably confounded in single time-point comparisons, making comparisons between studies difficult. By sampling populations before and after treatment and including untreated controls, our design allowed to distinguish seasonal demographic changes from the effect of fungicides. Both fully sampled fields lose low-frequency SNPs across the season while π stays buffered, the expected signature of a mild contraction in which low-frequency alleles are lost faster than mean pairwise nucleotide diversity (Nei *et al*., 1975) (Figure 3A, F). The rare tails erode differently in the two fields, and only at the spray step. In Kollmar the fungicide -treated population is singleton-poorer than its untreated control, deepening the seasonal depletion; in Dollerup it is singleton-richer, offsetting it (Supplementary Table 8b). The same compound, applied in the same week to the same cultivar, therefore moves the frequency spectrum in opposite directions in two fields within ∼ 117 Km. Thus, epidemic progression sets the demographic trajectory, whereas fungicide acts as a local perturbation whose magnitude and direction depend on the starting population. This interpretation is consistent with observations from natural epidemics, where early-established pathogen strains disproportionately shape the genetic composition of populations later in the epidemic, resulting in seasonal shifts in strain composition and epidemic trajectories (Eck *et al*., 2022) and theoretical models predict that seasonal epidemic dynamics alone can maintain pathogen diversity by promoting the coexistence of strains with different life-history strategies, highlighting seasonality as an important evolutionary force independent of spatial population structure (Andreasen & Dwyer, 2023). Here, we extend these ideas by showing that, at the genome-wide level, seasonal epidemic progression consistently outweighs the additional effect of fungicide application in shaping short-term population change.

### Selection acts on a shared standing variation reservoir

The fungicide response should thus be interpreted as selection acting on an already evolving population, rather than initiating adaptation de novo. Allele-frequency changes were observed genome-wide, and because these populations never evolve free of selection, sweeps are already present at the early baseline. The treatment contrast therefore tests the intensification of selection, not its presence (Croll & McDonald, 2017; Hartmann *et al*., 2021). Although both paired fields shared a common reservoir of standing variation, they differed in CYP51 composition and the abundance of rare variants before treatment. Consequently, identical fungicide applications acted on different evolutionary backgrounds, producing contrasting genomic responses despite the same management regime. Consistent with this, only a single CY P51 substitution (S524T) showed a treatment-specific increase, while all other major resistance alleles were already segregating before fungicide application. Together, these observations support the view that azole adaptation in European *Z. tritici* populations is driven primarily by changes in the frequencies of standing resistance alleles rather than by the repeated emergence of novel target-site mutations (Huf *et al*., 2018; Hellin *et al*., 2021; Kildea *et al*., 2025).

Effector genes showed the same overarching pattern. Rather than the gain or loss of major haplotypes, epidemic progression primarily reshaped the relative frequencies of existing effector haplotypes while maintaining overall diversity. This agrees with increasing evidence that natural *Z. tritici* populations harbour extensive standing effector diversity, including our previous observations from the UK population (Tobian Herreno et al., 2025), and that adaptation proceeds largely through changes in the freque ncies of pre-existing virulence alleles. This has been demonstrated for *Avr3D1, AvrStb9* and *AvrStb16q*, where virulence alleles were already present in field populations before selection favoured their increase (Meile et al., 2018; Amezrou et al., 2023; Orellana-Torrejon et al., 2022a). Similar patterns have recently been described in wheat powdery mildew, where standing diversity at the AvrPm17 locus allowed rapid adaptation following deployment of the Pm17 resistance gene, illustrating that selection on pre-existing effector diversity may be a common feature of rapidly evolving cereal pathogens (Müller *et al*., 2022). Together, these observations confirm that adaptation in *Z. tritici* predominantly exploits functional standing genetic variation and show that seasonal epidemic progression reshapes this variation while largely preserving the adaptive reservoir on which future adaptation depends.

### Implications: durability is a population-genetics problem

These dynamics recast the durability of both fungicides and resistance genes as a property of the pathogen population rather than of the control measure itself. Because Zt combines a very large effective population size with frequent recombination, adaptation is unlikely to be limited by the appearance of new mutations but instead by selection acting on standing variation.(Bernasconi *et al*., 2022; McDonald *et al*., 2022; Amezrou *et al*., 2024). Consequently, every new fungicide or resistance gene acts on a population that already contains substantial adaptive potential, helping explain why even pyramided resistance genes might not be durable in *Z. tritici* (Stam & McDonald, 2018; Saintenac *et al*., 2021; Battache *et al*., 2022, 2024; Suffert *et al*., 2024; Meile *et al*., 2024).

Two practical implications follow. First, because adaptation proceeds from standing variation that differs among fields, local population composition should increasingly be considered when designing disease management strategies (Talas *et al*., 2026). Second, adaptive alleles emerge from the low-frequency tail of the allele-frequency spectrum, surveillance should prioritize sufficiently dense sampling to detect these variants before they rise in frequency and importantly, sample deeply and densely enough to resolve variation below conventional filtering thresholds (Harrison *et al*., 2024). The breakdown of *Stb16q* in France exemplifies this process: resistance failed through the expansion of pre-existing virulence alleles rather than the appearance of novel mutations, illustrating the value of genomic surveillance focused on standing variation (Saintenac *et al*., 2021; Orellana-Torrejon *et al*., 2022a; Battache *et al*., 2022; Suffert *et al*., 2024), yet, the emergence that dense genomic monitoring can already track across fields (Harrison *et al*., 2024; Talas *et al*., 2026). Together, these findings highlight standing genetic variation as the principal substrate of short-term adaptation and identify its monitoring and management as a central challenge for improving the durability of both fungicides and host resistance.

## Acknowledgements

We want to thank Susanne Kleingarn for her assistance in the isolate preparation and culturing, Dr. Holger Klink for the coordination of the field samplings and Dr. Andreas Buechse for the sampling design.

## Supplementary figures

**Supplementary Figure S1.**
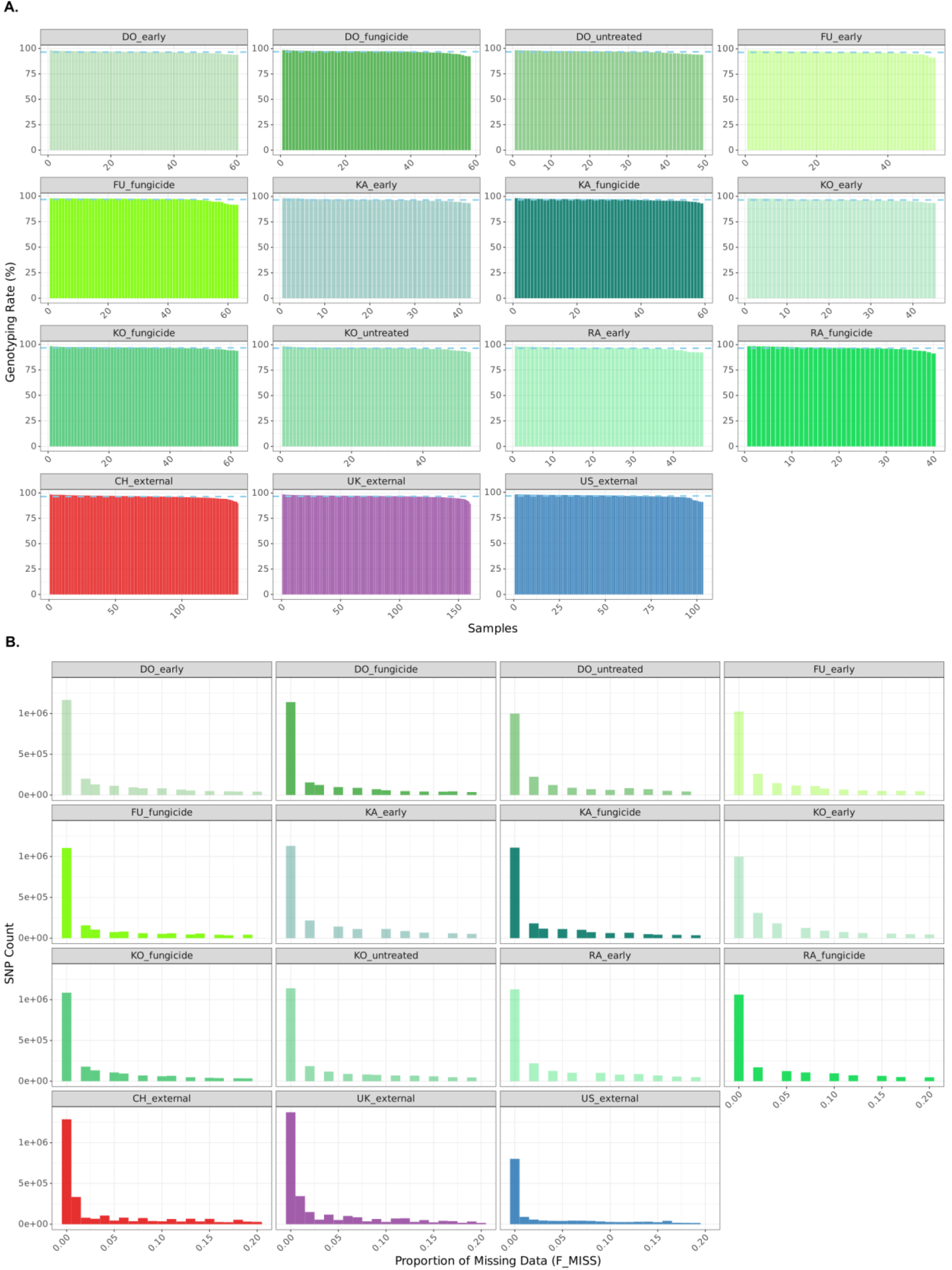
Genotyping was high quality and comparable across all eight fields after filtering. **A.** Per-isolate genotyping rate for the UK, US, CH and German fields, from PLINK missingness reports. **B.** Per-site missingness for the same fields.

**Supplementary Figure S2.**
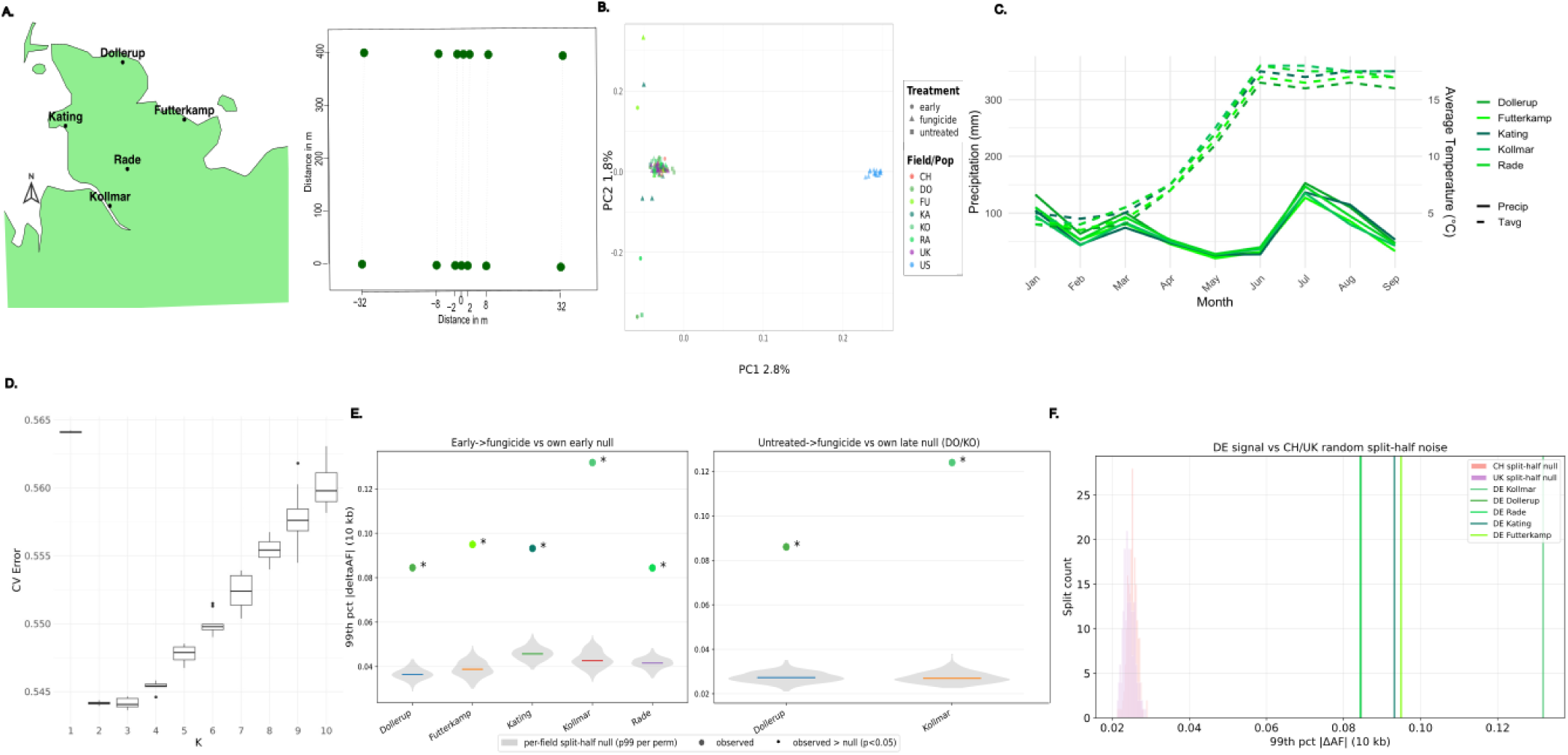
Sampling design, field-level population structure, climate, and the null models for allele-frequency change. **A.** Field locations in Schleswig-Holstein and the sampling layout: two exponential transects per field in the outermost lanes, seven points each (a centre point plus 2, 8 and 32 m to either side). **B.** PCA of the field-representative isolates. **C.** Monthly precipitation and the min-max temperature range in the collection year, coloured per field. Temperature range did not differ (Kruskal-Wallis P > 0.05); precipitation differed: Futterkamp and Kollmar each differed from Dollerup and Rade (Wilcoxon rank-sum, Benjamini-Hochberg, p < 0.05), while Kating was intermediate and differed from none. **D.** Cross-validation error across K for the admixture analysis (core SNPs, MAF 5%, thinned to one SNP per 10 kb). **E.** Observed 99th percentile of the absolute ΔAF in 10 kb windows for each between-time-point and between-treatment contrast, compared with a per-field split-half null obtained by randomly halving the early population, testing whether the allele-frequency change exceeds within-field sampling noise. **F.** Observed 99th percentile of the absolute ΔAF for the German between-time-point contrasts compared with split-half null distributions from randomly subsampled Swiss and United Kingdom isolates, testing whether the German signal exceeds the noise produced by comparisons within single-time-point European fields.

**Supplementary Figure S3.**
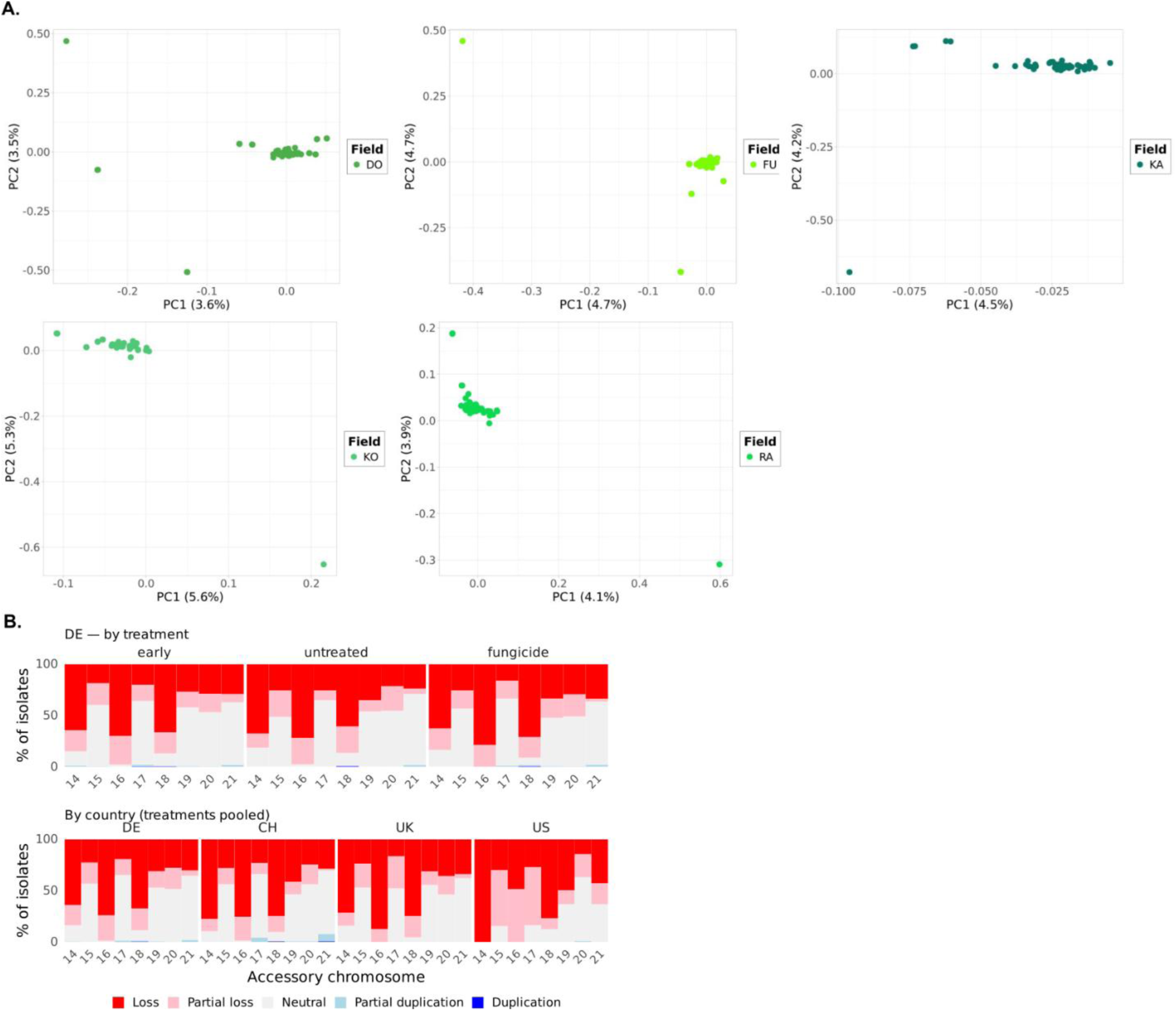
Within-field structure is weak and heterogeneous: no field-level PCA separation, but diversity and accessory copy number differ between fields. **A.** PCA analyses per field at the early time point were done using the core genome SNPs of each field with the SNPrelate package. **B.** Accessory-chromosome (14–21) copy-number variation across the whole dataset, called with CNVkit (v 0.9.10), the coverage ratio per chromosome was classified as loss, partial loss, neutral, partial duplication or duplication, and bars show the percentage of isolates in each class. The upper panel shows the German fields broken down by treatment (early, untreated, fungicide); the lower panel shows the four countries (Germany, Switzerland, the United Kingdom, United States) with treatments pooled. Per-chromosome copy number (mean log2 ratio) was compared between treatments within each field, between German fields, and between countries using Kruskal–Wallis tests (rank-sum for two-group comparisons), with an additional two-sided Wilcoxon rank-sum test for differences between the untreated and fungicide treatment; all were Benjamini–Hochberg corrected across the eight accessory chromosomes within each comparison.

**Supplementary Figure S4.**
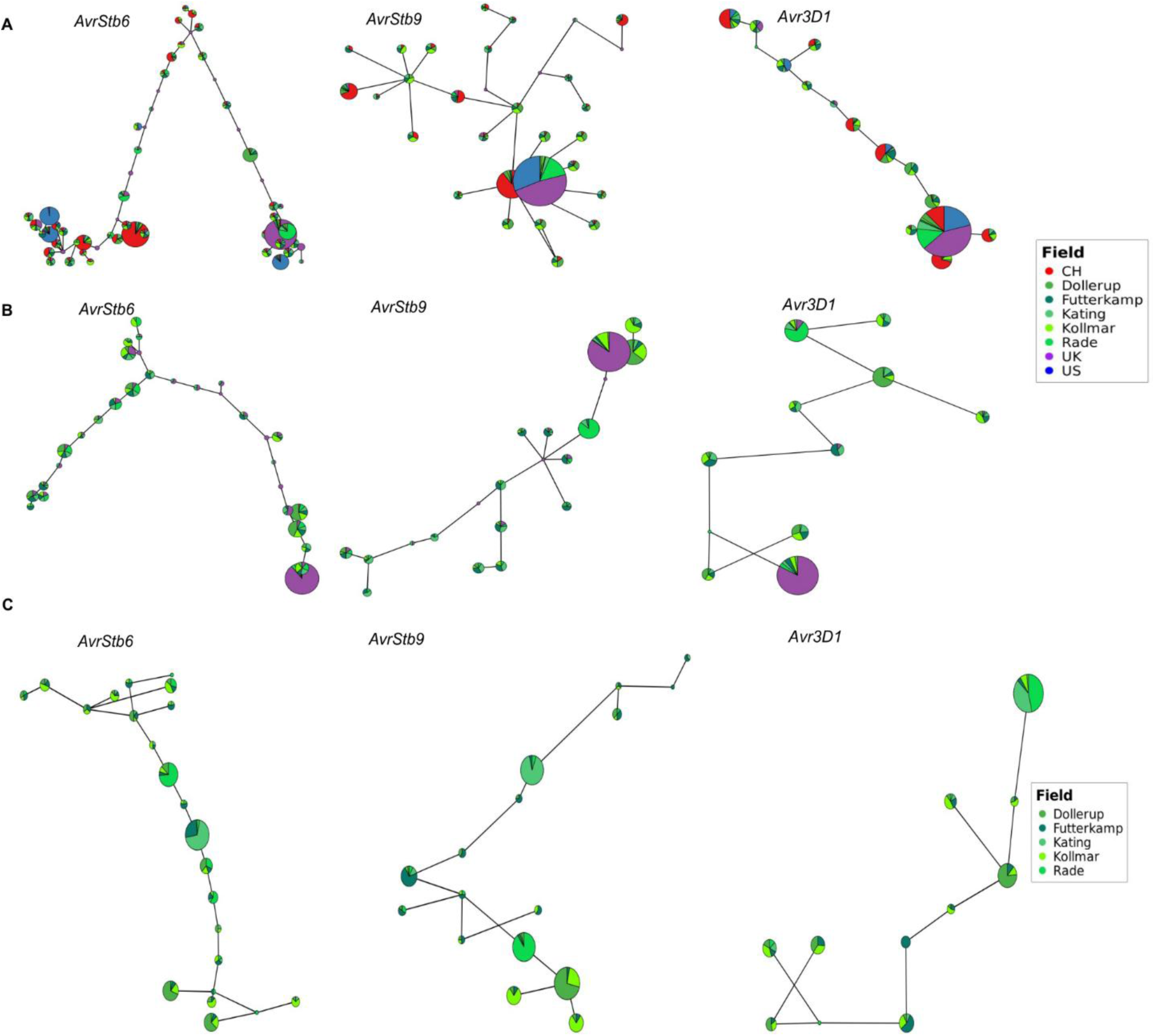
Characterised-effector haplotypes are regionally shared: no private multilocus genotypes separate the fields. **A.** All fields. **B.** European fields. **C.** German fields.

**Supplementary Figure S5.**
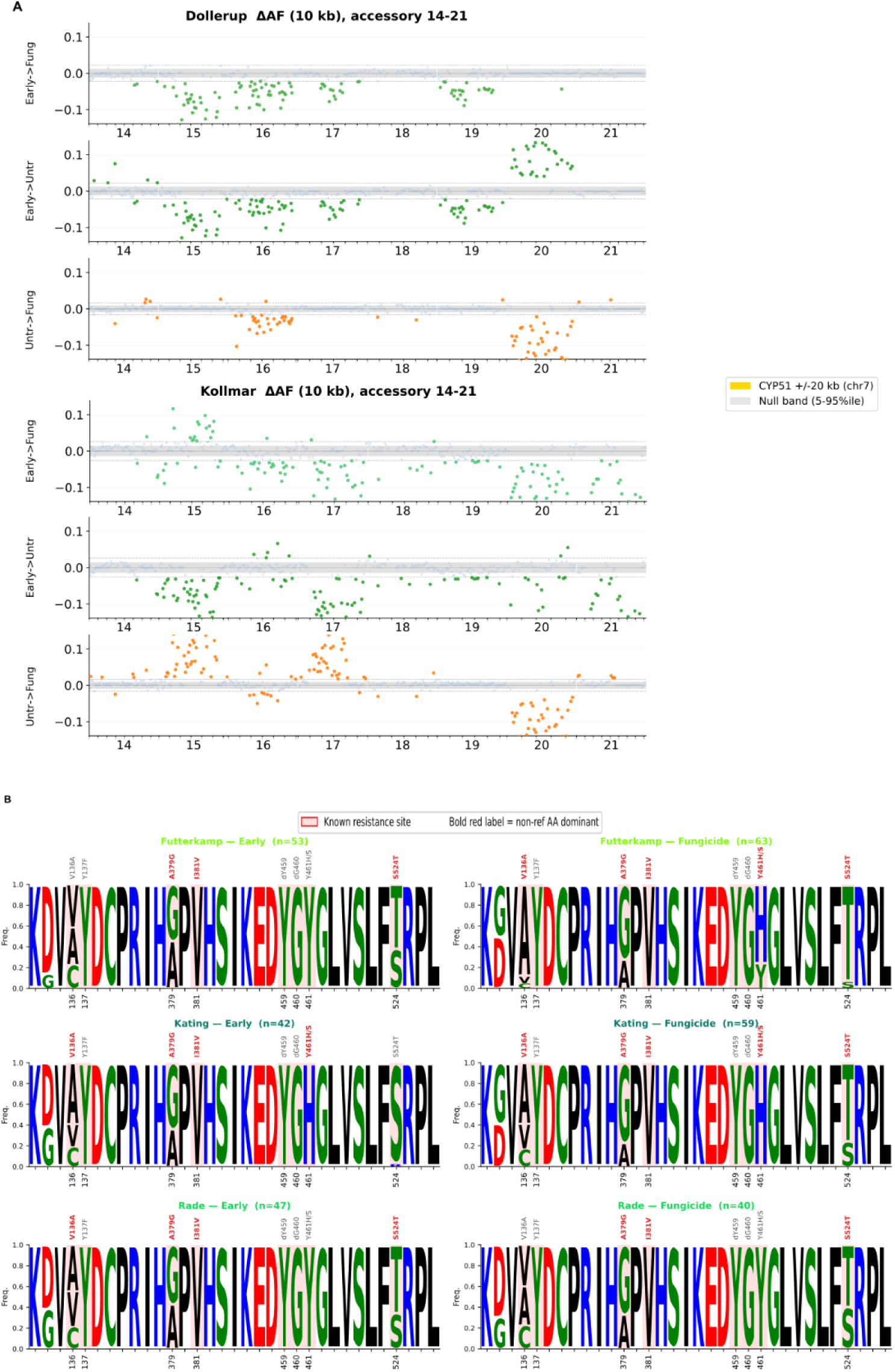
Allele-frequency change extends across the accessory genome, and CYP51 resistance substitutions are dominant before treatment in every field. With the clarification that presence-absence variation is confounded in the Accessory PCA with actual ΔAF SNP shift **A.** Mean ΔAF across the accessory genome between treatments, early to untreated and untreated to fungicide. **B.** CYP51 motif plots for all German fields and treatments; literature-evidenced resistance sites are boxed in red, and a bold red label marks positions where the non -reference allele dominates.

**Supplementary Figure S6.**
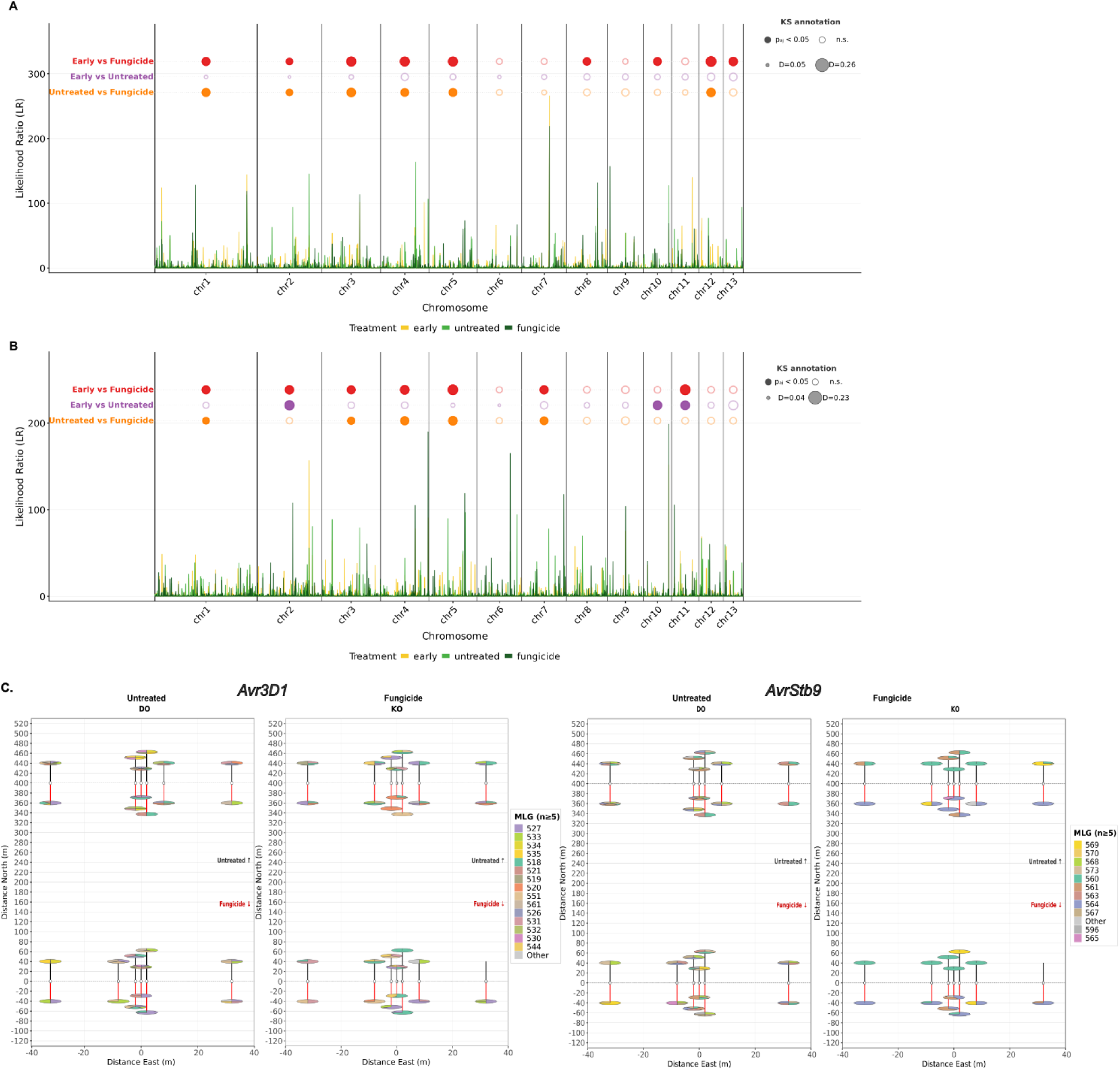
Selective sweeps are present at the early baseline and shift, not appear, under treatment; effector haplotypes coexist within single plants. **A.** Core-genome sweeps across treatments in Dollerup, with distribution differences by Kolmogorov-Smirnov (P < 0.05) and effect size D. **B.** The same for Kollmar. **C.** Per-plant MLG composition for Dollerup and Kollmar across the sampling points, one pie chart per plant, for both late - season treatments for *AvrStb9 and Avr3D1*.

